# The S1/S2 boundary of SARS-CoV-2 spike protein modulates cell entry pathways and transmission

**DOI:** 10.1101/2020.08.25.266775

**Authors:** Yunkai Zhu, Fei Feng, Gaowei Hu, Yuyan Wang, Yin Yu, Yuanfei Zhu, Wei Xu, Xia Cai, Zhiping Sun, Wendong Han, Rong Ye, Hongjun Chen, Qiang Ding, Qiliang Cai, Di Qu, Youhua Xie, Zhenghong Yuan, Rong Zhang

## Abstract

The global spread of SARS-CoV-2 is posing major public health challenges. One unique feature of SARS-CoV-2 spike protein is the insertion of multi-basic residues at the S1/S2 subunit cleavage site, the function of which remains uncertain. We found that the virus with intact spike (Sfull) preferentially enters cells via fusion at the plasma membrane, whereas a clone (Sdel) with deletion disrupting the multi-basic S1/S2 site instead utilizes a less efficient endosomal entry pathway. This idea was supported by the identification of a suite of endosomal entry factors specific to Sdel virus by a genome-wide CRISPR-Cas9 screen. A panel of host factors regulating the surface expression of ACE2 was identified for both viruses. Using a hamster model, animal-to-animal transmission with the Sdel virus was almost completely abrogated, unlike with Sfull. These findings highlight the critical role of the S1/S2 boundary of the SARS-CoV-2 spike protein in modulating virus entry and transmission.

## INTRODUCTION

SARS-CoV-2 and SARS-CoV share nearly 80% nucleotide sequence identity and use the same cellular receptor, angiotensin-converting enzyme 2 (ACE2), to enter target cells(Hoffmann et al., 2020b; Zhou et al., 2020). However, the newly emerged SARS-CoV-2 exhibits greater transmissibility(Cespedes and Souza, 2020; Chen, 2020; Hui et al., 2020; Li et al., 2020). The viral structural protein, spike (S), plays critical roles in determining the entry events, host tropism, pathogenicity, and transmissibility. One significant difference between the SARS-CoV-2 spike protein and those of other bat-like SARS-CoV is the insertion of multi-basic residues (RRAR) at the junction of S1 and S2 cleavage site(Wang et al., 2020b). Previous studies showed that expression of SARS-CoV-2 spike in cells promotes cell-cell membrane fusion, which is reduced after deletion of the RRAR sequence or when expressing SARS-CoV S protein lacking these residues(Hoffmann et al., 2020a; Xia et al., 2020). Pseudovirus or live virus bearing SARS-CoV-2 spike deletion at the S1/S2 junction decreased the infection in Calu-3 cells and attenuated infection in hamsters(Hoffmann et al., 2020a; Lau et al., 2020). The sequence at the S1/S2 boundary seems to be unstable, as deletion variants are observed both in cell culture and in patient samples(Lau et al., 2020; Liu et al., 2020; Ogando et al., 2020; Wong et al.). SARS-CoV-2 entry is mediated by sequential cleavage at the S1/S2 junction site and additional downstream S2’ site of spike protein. The sequence at the S1/S2 boundary contains a cleavage site for the furin protease, which could preactivate the S protein for membrane fusion and potentially reduce the dependence of SARS-CoV-2 on plasma membrane proteases, such as transmembrane serine protease 2 (TMPRSS2), to enable efficient cell entry(Shang et al., 2020). Here, we evaluate how the deletion at the S1/S2 junction impacts virus entry and cell tropism, define the host factors regulating this process, and determine whether the presence of these multi-basic residues contributes to the enhanced transmission of SARS-CoV-2.

## RESULTS

### The deletion at the S1/S2 boundary of spike protein impacts the infectivity in cells

We observed the same phenomena that others have reported, an instability of the SARS-CoV-2 S1/S2 boundary(Lau et al., 2020; Liu et al., 2020; Ogando et al., 2020). Using the patient-isolated SARS-CoV-2 SH01 strain, we performed three rounds of plaque purification in Vero E6 cells in the presence of trypsin and observed no mutations in any of the structural genes (Sfull virus). However, after two additional rounds of passage without trypsin, a 21-nucleotide deletion at the S1/S2 cleavage site was acquired, disrupting the RRAR motif (**Figure 1A, Figure S1A**). We designated the plaque-purified deletion clone as Sdel virus and detected no additional mutations in the full-length genome when compared to the Sfull virus. Unexpectedly, this presumed cell culture adaptation could be prevented by adding trypsin to the media or by ectopically expressing the serine protease TMPRSS2 in Vero E6 cells (**Figure S1B and 1C**). Compared to Sfull, the deletion-bearing Sdel virus exhibited a dramatic increase in infectivity as measured by the greater percentage of nucleocapsid (N) antigen-positive cells (**Figure S2A**) and higher yield in virus production in wild-type Vero E6 (hereafter Vero cells), Vero plus trypsin, Vero expressing TMPRSS2, and A549 cells expressing the receptor ACE2 (**Figure 1B-D**). Conversely, in human Calu-3 lung epithelial cells, the Sdel virus replicated slower than the Sfull clone (**Figure 1B-D**), similar to previous reports using a pseudovirus or fully infectious, mutant virus (Hoffmann et al., 2020a; Lau et al., 2020). Moreover, we found that pseudovirus bearing the S protein from Sfull, Sdel, or a RRAR mutant variant (R682S, R685S)(Wang et al., 2020a), had a phenotype similar to infectious viruses used in these cell types (**Figure 1D and 1E**). Of note, infection using either the Sdel S- or mutant variant S (R682S, R685S)-bearing pseudovirus was decreased by approximately ten-fold in Calu-3 cells, highlighting the critical role of these basic residues at the S1/S2 boundary in infectivity.

**Figure 1.**
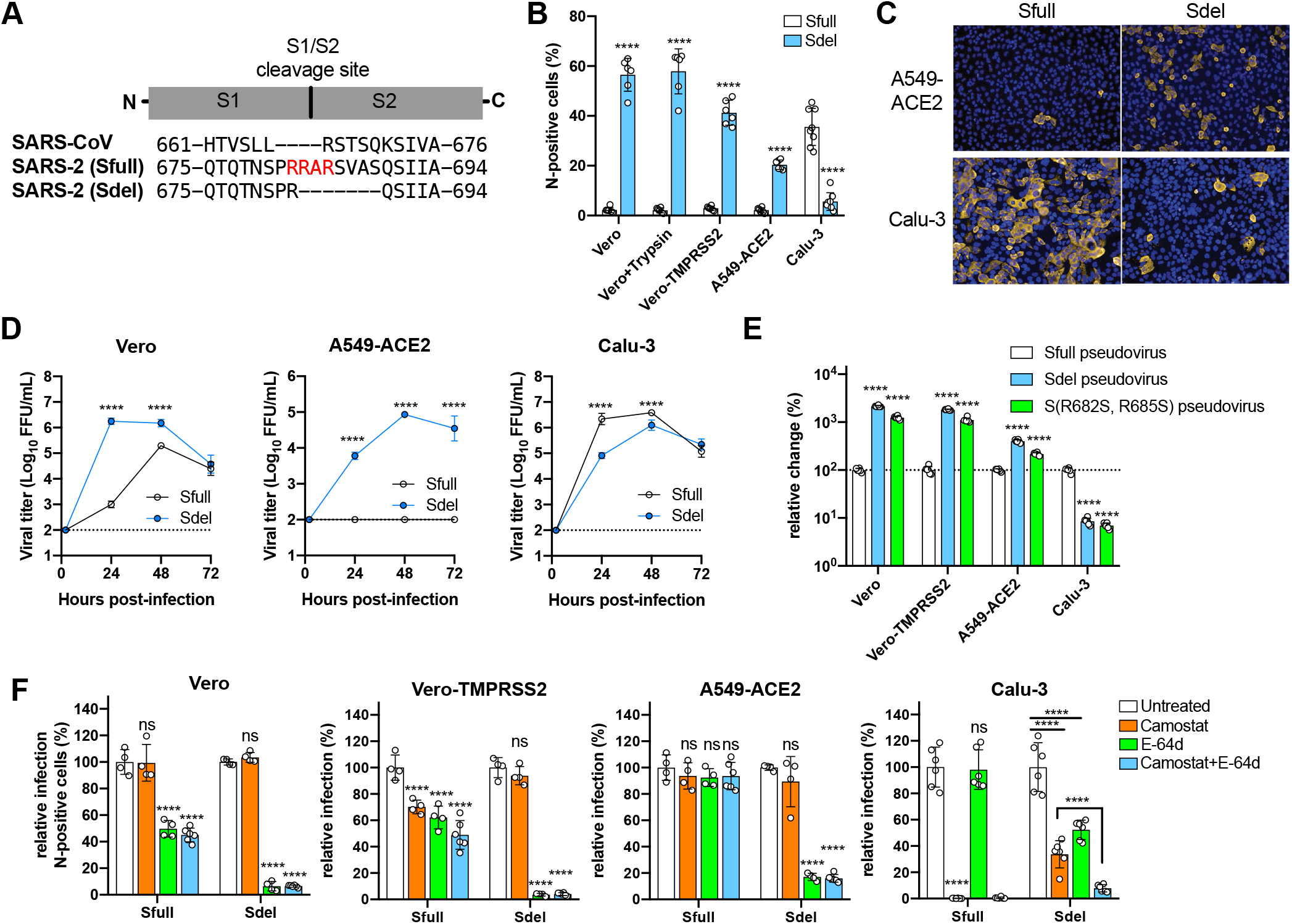
SARS-CoV-2 bearing the deletion at the S1/S2 junction site of spike protein preferentially enters cells through the endosomal pathway. **A.** Sequence alignment of spike protein encompassing the cleavage site between S1 and S2 subunits. The spike proteins of SARS-CoV-2 without (Sfull strain) and with (Sdel strain) deletion were used to compare with that of SARS-CoV. The insertion of multi-basic amino acids in spike protein of SARS-CoV-2 was shown in red. **B.** Comparison of the replication property between Sfull and Sdel strains in different cell lines. The percentage of nucleocapsid (N) protein positive cells was analyzed by imaging-based analysis following virus infection. Data shown are an average of two independent experiments performed in triplicate. **C.** The immunofluorescence staining of N protein in A549-ACE2 and Calu-3 cells infected with Sfull or Sdel virus. A representative of two independent experiments was shown. **D.** Assessment of live virus production in different cell lines infected with Sfull or Sdel strain. Data are pooled from two independent experiments conducted in triplicate. **E.** Evaluation of entry efficiency in different cell lines infected with pseudoviruses bearing spike protein Sfull, Sdel, or S mutant (R682S, R685S). Data shown are an average of two independent experiments performed in triplicate and are normalized to the Sfull of individual experiments. **F.** Effect of TMPRSS2 serine protease inhibitor Camostat and cysteine protease inhibitor E-64d on Sfull or Sdel infection in different cell lines. Data shown are an average of two independent experiments performed in duplicate or triplicate and are normalized to the untreated group of individual experiments. One-way ANOVA with Dunnett’s test (A, B, C); two-way ANOVA with Sidak’s test (B); ****P < 0.0001; ns, not significant.

### S1/S2 boundary of spike protein modulates cell entry pathways

Coronavirus enters cells through two pathways: fusion at the plasma membrane or in the endosome(Tang et al., 2020). To assess the impact of the S1/S2 junction deletion on viral entry, cells were treated with camostat mesilate, a TMPRSS2 inhibitor that blocks viral fusion at the plasma membrane, and/or E-64d (aloxistatin), an inhibitor that blocks the protease activity of cathepsins B and L, which are required for the endosomal membrane fusion (**Figure 1F, Figure S2B**). We observed apparent S1/S2 cleavage for Sfull virus but not for Sdel in multiple cell types (**Figure S3A**). Sfull virus infection, as measured by N antigen-positive cells, was sensitive to inhibition by E-64d but not camostat in Vero cells (**Figure 1F**). When TMPRSS2 was expressed, both camostat and E-64d inhibited the infectivity of Sfull, indicating that expression of TMPRSS2 could promote the membrane fusion entry pathway. Remarkably, E-64d and camostat had no effect on Sfull virus in A549-ACE2 cells, suggesting that in this cell Sfull may use other TMPRSS2 homologs or trypsin-like proteases to activate fusion at the plasma membrane since TMPRSS2 expression is absent in A549 cells(Matsuyama et al., 2020). We observed a similar phenotype even when cells were treated with a high concentration of inhibitors (**Figure S2C**). In Calu-3 cells, camostat completely blocked the Sfull infection, but E-64d had minimal effects, suggesting that Sfull preferentially enters Calu-3 cells via the plasma membrane fusion pathway.

For the Sdel virus, E-64d significantly inhibited infection in Vero, Vero-TMPRSS2, and A549-ACE2 cells, whereas camostat did not reduce the infection, even in Vero-TMPRSS2 cells (**Figure 1F**). It is noteworthy that Sdel was sensitive to both inhibitors in Calu-3 cells unlike the Sfull virus, and these two compounds exerted a synergetic effect on Sdel infection. This suggests Sdel utilizes both plasma membrane and endosomal fusion pathways in Calu-3 cells. The spike protein of SARS-CoV does not have the insertion of multiple basic residues at the S1/S2 cleavage site and thus resembles the Sdel virus (**Figure 1A**). Indeed, E-64, but not camostat, efficiently inhibited SARS-CoV pseudovirus infection in multiple cell types (**Figure S2D**). These results demonstrate that the deletion at the S1/S2 junction site propels the virus to enter cells through the endosomal fusion pathway, which is less efficient than the fusion pathway at the plasma membrane in airway epithelial cells as indicated by the reduced infectivity in Calu-3 cells. Both Sdel and SARS-CoV may share a similar entry pathway.

### CRISPR/Cas9 screen identifies endosomal entry factors required for Sdel virus infection

Genome-wide CRISPR/Cas9 screens have enabled the identification of host factors required for efficient virus infection(Karakus et al., 2019; Marceau et al., 2016; Richardson et al., 2018; Zhang et al., 2018; Zhang et al., 2016). A lack of suitable human physiologically relevant cell lines and the S protein-induced syncytia formation in cells have made such a screen for SARS-CoV-2 very challenging. We found that Sdel virus preferentially enters A549-ACE2 cells via the endosomal fusion pathway, replicates robustly, does not cause syncytia, and efficiently results in cell death. Because of these properties, we performed a genome-wide, cell survival-based screen with the Sdel virus in A549-ACE2 cells transduced with a library of single-guide RNAs (sgRNAs) targeting 19,114 human genes (**Figure 2A**)(Doench et al., 2016). The vast majority of transduced cells inoculated with Sdel virus died within seven days of infection. Surviving cells were harvested and expanded for a second round of challenge with Sdel. The remaining surviving cells were expanded and subjected to genomic DNA extraction, sgRNA sequencing, and data analysis (**Supplementary Tables 1 and 2**).

**Figure 2.**
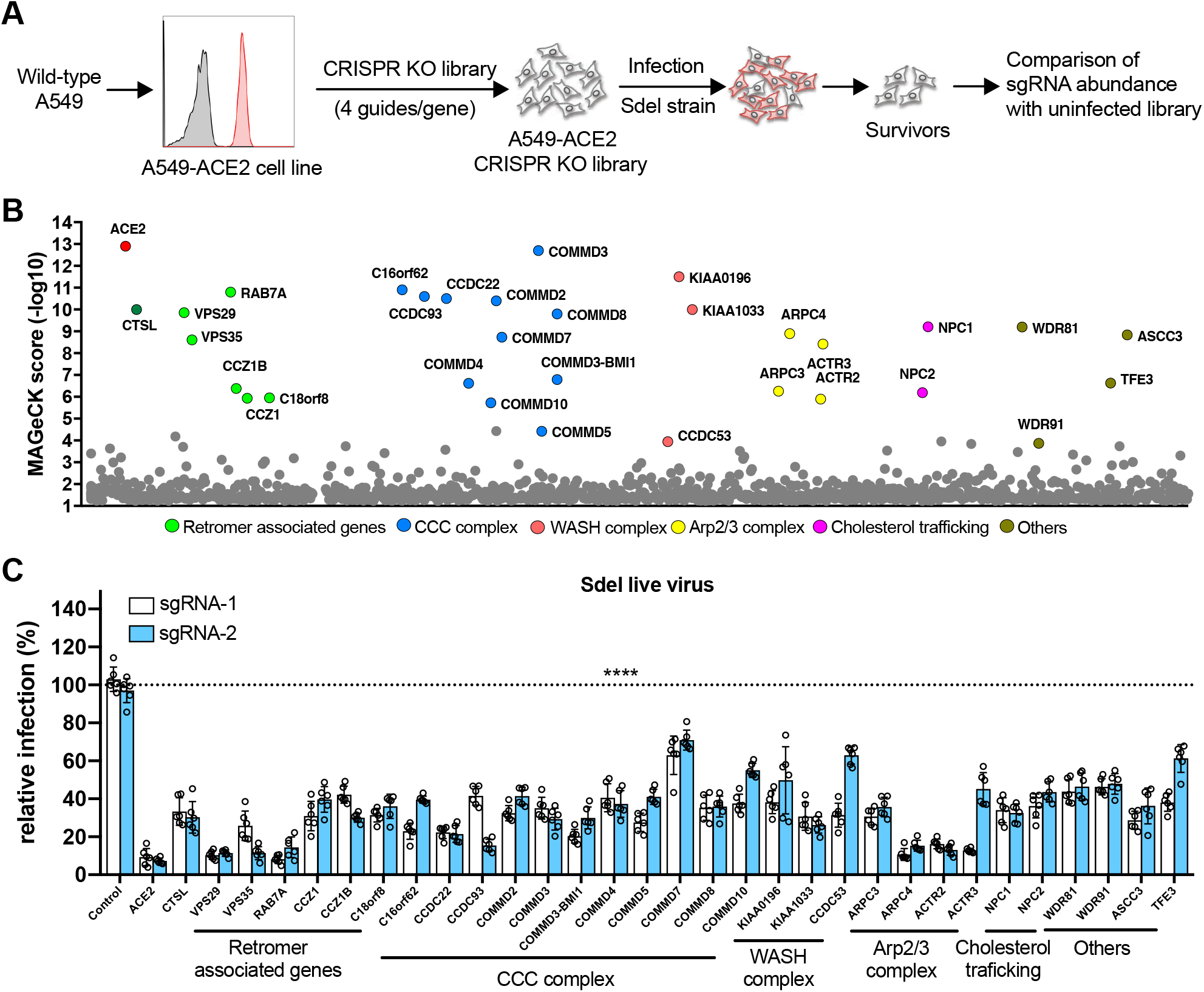
Genome-wide CRISPR/Cas9 screen identifies genes and pathways required for SARS-CoV-2 infection. **A.** Schematic of the screening process. A549 cells expressing the human ACE2 were used to generate the CRISPR sgRNA knockout cell library. The library was infected with Sdel strain of SARS-CoV-2, and cells survived were harvested for genomic extraction and sequence analysis. **B.** Genes and complexes identified from the CRISPR screen. Of the top 36 hits (FDR<0.15), the validated genes (from Figure 2C) were indicated based the MAGeCK score. **C.** Top 36 genes were selected for experimental validation in A549-ACE2 cells using two independent sgRNAs by Sdel live virus infection, and only the 32 genes that showed statistic significance were indicated. Data shown are an average of two independent experiments performed in triplicate and are normalized to the controls of individual experiments. One-way ANOVA with Dunnett’s test; ****P < 0.0001.

The top candidates from the CRISPR screen were determined according to their MAGeCK score (**Figure 2B**). The top hit was ACE2, the cellular receptor that confers susceptibility to SARS-CoV-2, which confirmed the validity of the screen. Additionally, the gene encoding cathepsin L (CTSL), a target of our earlier assay using E-64d that is known to be important for activating SARS-CoV virion membrane fusion with the endosome (Simmons et al., 2005), also was identified, again confirming the utility of the screening strategy.

We chose the top 36 genes with a cutoff of false discovery rate (FDR) < 0.15. For each specific gene target, A549-ACE2 cells were transduced with two independent sgRNAs and then infected with Sdel. The percentage of N protein-positive cells was determined by image-based analysis. Remarkably, editing of all 32 genes resulted in a statistically significant reduction in Sdel infection compared to cells receiving the control sgRNA (**Figure 2C**). Most of these genes were associated with the endolysosome, including components of the retromer complex, the COMMD/CCDC22/CCDC93 (CCC) complex, Wiskott-Aldrich syndrome protein and SCAR homologue (WASH) complex, and actin-related protein 2/3 (Arp2/3) complex, which have significant roles in endosomal cargo sorting(Liu et al., 2016; McNally and Cullen, 2018). We also identified genes encoding the WD Repeat Domain 81 (WDR81)-WDR91 complex, which was detected in a previous a genetic screen for regulators of endocytosis and the fusion of endolysosomal compartments(Rapiteanu et al., 2016). Similarly, we identified the gene encoding Transcription Factor Binding To IGHM Enhancer 3 (TFE3), which may regulate lysosomal positioning in response to starvation or cholesterol-induced lysosomal stress(Willett et al., 2017). We also validated NPC Intracellular Cholesterol Transporter 1 (NPC1) and NPC2, which regulate intracellular cholesterol trafficking, as important for Sdel infection(Cologna and Rosenhouse-Dantsker, 2019; Pfeffer, 2019). In addition, the gene for Activating Signal Cointegrator complex 3 (ASCC3), which functions as a negative regulator of the host defense response, was identified in our screen(Li et al., 2013). From these hits, we selected representative genes to validate for cell-type specificity in HeLa-ACE2 cells, finding that all the genes tested greatly reduced infection with Sdel virus (**Figure S4A**).

To define the stage of viral infection that each of the 32 validated genes acted, one representative sgRNA per gene was selected for study in A549-ACE2 cells. Due to its known antiviral activity, *ASCC3* was not targeted. We confirmed that editing of these genes did not affect cell viability (**Figure S4B**). The gene-edited cells were infected with pseudovirus bearing the Sdel virus S protein or, as a control, the glycoprotein of vesicular stomatitis virus (VSV-G) (**Figure 3A and 3B**). Consistent with data from the fully infectious Sdel virus, editing any of the selected genes markedly inhibited Sdel pseudovirus infection whereas only editing of some retromer-associated genes and the Arp2/3 complex significantly reduced the VSV-G pseudovirus infection. These results suggest that these genes mediate Sdel virus entry. Notably, pseudovirus bearing the spike protein of SARS-CoV, which lacks the multiple basic residues at the S1/S2 junction as Sdel, exhibited a phenotype similar to Sdel pseudovirus and Sdel live virus (**Figure 3C and 2C**). Editing of these genes, including those encoding CTSL, cholesterol transporters NPC1/2, WDR81/91, and TFE3, markedly reduced infection, suggesting that Sdel and SARS-CoV may utilize similar entry machinery (**Figure 3C**). Intriguingly, these genes edited also significantly inhibited the infection by pseudovirus bearing the spike protein of MERS-CoV in A549-ACE2-DPP4 cells (**Figure 3D**). Although the furin cleavage site is present at the S1/S2 boundary of MERS-CoV{Millet, 2014 #374}, it preferentially enters the A549 cell via endosomal pathway as indicated by its sensitivity to E-64d inhibitor (**Figure S2E**). This is possibly due to the lack of proper protease to activate the plasma membrane fusion pathway in A549 cells for MERS-CoV as compared to the Sfull virus.

**Figure 3.**
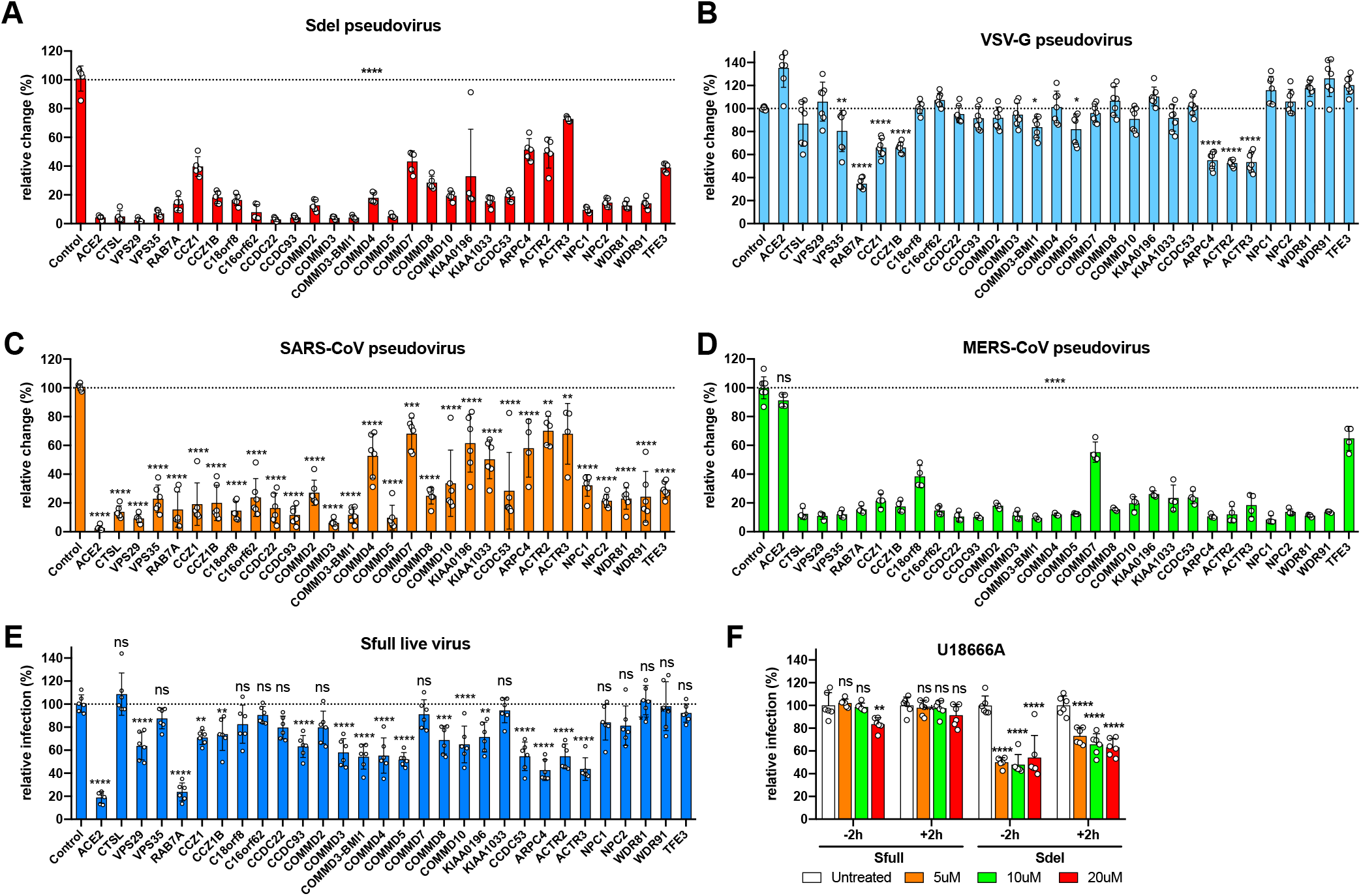
Genes identified are required for the cell entry of SARS-CoV-2, SARS-CoV, and MERS-CoV. **A-D.** The genes (selected from Figure 2C) were verified for the infection by pseudovirus bearing the spike protein of SARS-CoV-2 Sdel strain (A), the the glycoprotein of vesicular stomatitis virus (VSV-G) (B), the spike protein of SARS-CoV (C), or the spike protein of MERS-CoV (D). One representative sgRNA per gene was used in A549-ACE2 cells. **E.** The genes selected were verified for the infection by the SARS-CoV-2 Sfull live virus. **F.** Effect of NPC1 inhibitor U18666A on virus infection. Cells were treated with U18666A at the indicated concentrations 2 h prior to or 2 h post infection by Sfull or Sdel live virus. The viral N-positive cells were calculated. Data shown are an average of two independent experiments performed in duplicate or triplicate and are normalized to the controls of individual experiments. One-way ANOVA with Dunnett’s test; **P < 0.01; ***, P < 0.001; ****P < 0.0001; ns, not significant.

### Sdel endosomal entry factors are not required for Sfull virus infection

To determine whether these genes identified impact Sfull virus infection, one representative sgRNA per gene was tested (**Figure 3E**). The editing efficiency of some these genes by sgRNAs was confirmed by western blotting (**Figure S3B**). As expected, editing of *CTSL* did not reduce infection, as the Sfull virus enters A549-ACE2 cells via an endosomal-independent pathway (demonstrated in **Figure 1F**). In general, editing of genes encoding complexes that regulate the retrieval and recycling of cargo significantly reduced infection, albeit to a lesser extent than observed with the Sdel live virus. However, unlike our results with the Sdel virus, editing of *NPC1* or *NPC2* had a negligible impact on Sfull virus infection, raising question of the effectiveness of perturbing cholesterol trafficking with inhibitors such as U18666A in COVID-19 as previously proposed(Ballout et al., 2020; Sturley et al., 2020).

U18666A, a cationic sterol, binds to the NPC1 protein to inhibit cholesterol export from the lysosome, resulting in impaired endosome trafficking, late endosome/lysosome membrane fusion(Cenedella, 2009; Ko et al., 2001; Lu et al., 2015). U1866A has been shown to inhibit the S protein-driven entry of SARS-CoV, Middle East Respiratory Syndrome coronavirus (MERS-CoV), and the human coronaviruses NL63 and 229E, with the most efficient inhibition observed with SARS-CoV(Wrensch et al., 2014). The antiviral effect of U18666A on type I feline coronavirus (FCoV) has also been characterized *in vitro* and *in vivo(Doki et al., 2020; Takano et al., 2017)*. We found that, pretreating A549-ACE2 cells 2 h prior to or post infection had no inhibitory effect on Sfull virus (**Figure 3F**). In contrast, Sdel virus was more sensitive to U18666A, even when used for treatment 2 h post infection, presumably due to Sdel preferential usage of the endosomal entry pathway. Together with results showing no impact on Sfull virus infection after editing of genes *WDR81/91* and *TFE3* functioning in endolysosomes (**Figure 3E**), these studies suggest that, because of the different entry pathways used by the virus depending on the deletion at the S1/S2 boundary, the Sdel CRISPR hits in the endosomal pathway are dispensable for Sfull virus infectivity, and targeting the endosomal entry pathway with hibitors might be not efficient to block the virus infection.

### Genes regulating the ACE2 surface expression are required for infection by both Sdel and Sfull viruses

The Sdel-validated genes that also affected Sfull infectivity were largely multi-protein complexes (**Figure 2C and 3C**). These complexes are important for maintaining plasma membrane and lysosomal homeostasis by maintaining expression of key integral proteins, including signaling receptors and transporters(McMillan et al., 2017; McNally and Cullen, 2018). We hypothesized that disruption of these complexes might affect the binding or transit of virions. To this end, we performed binding and internalization assays using Sfull virus in A549-ACE2 cells. The genes *COMMD3, VPS29*, and *CCDC53*, which encode proteins that are comprise CCC, retromer, and WASH complexes, respectively, were each edited; effects on expression were confirmed by western blotting (**Figure S3B**). Notably, binding and internalization of Sfull virions to these cells was significantly decreased compared to control sgRNA (**Figure 4A**).

**Figure 4.**
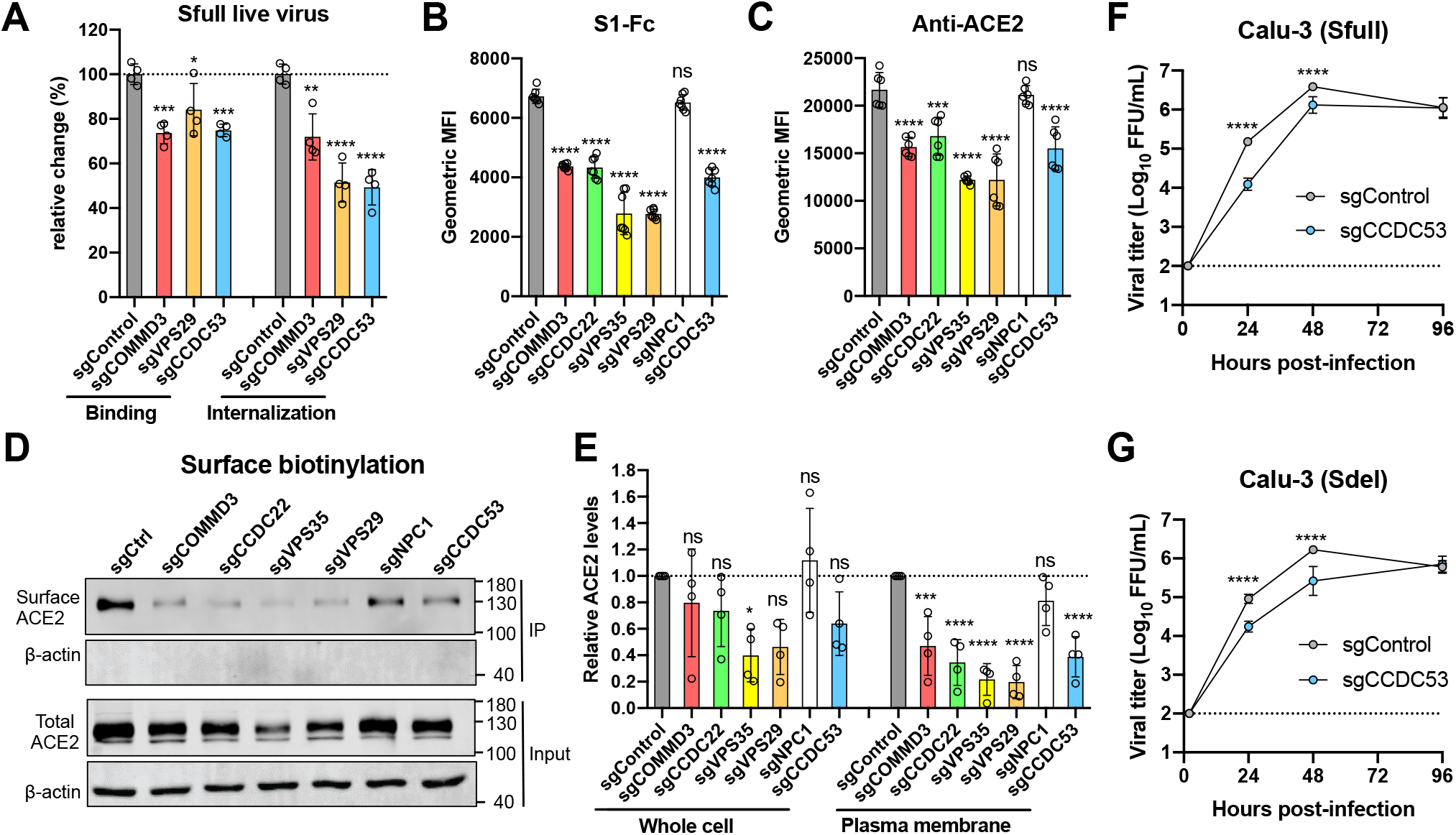
Genes identified are required for virus entry by regulating the expression of receptor ACE2. **A.** The effect on virion binding and internalization in gene-edited cells. A549-ACE2 cells were incubated with SARS-CoV-2 Sfull infectious virus on ice for binding or then switched to 37°C for internalization. Viral RNA was extracted for RT-qPCR analysis. Data shown are an average of two independent experiments performed in duplicate and are normalized to the controls of individual experiments. **B-C.** Surface expression of receptor ACE2 was decreased in gene-edited cells as measured by flow cytometry using S1-Fc recombinant protein or anti-ACE2 antibody. **D-E.** Surface and total expression of receptor ACE2 were decreased in gene-edited cells. The plasma membrane proteins were biotin-labeled and immunoprecipitated by streptavidin beads for Western blotting. One representative blot was shown (D) and Data are pooled from four independent experiments, quantified, and normalized to the controls of individual experiments (E). One-way ANOVA with Dunnett’s test (A, B, C, E); *P < 0.05; **P < 0.01; ***, P < 0.001; ****P < 0.0001; ns, not significant. **F-G.** The impact on viral production in CCDC53 gene-edited Calu-3 cells. The gene CCDC53 with significant reduction in virion internalization in Figure 4 (A) was selected for CRISPR sgRNA editing. The mixed cell population was infected with Sfull (F) or Sdel (G) to assess the virus yield. Two-way ANOVA with Sidak’s test; ****P < 0.0001.

The entry receptor ACE2 is critical for SARS-CoV-2 infection. To determine whether cell surface expression of ACE2 is regulated by these complexes, gene-edited cells *(COMMD3, VPS29, VPS35, CCDC53, CCDC22, and NPC1)* were incubated with S1-Fc recombinant protein or an anti-ACE2 antibody, and binding was measured by flow cytometry (**Figure 4B and 4C**). Editing of these genes perturbed the surface expression of ACE2, with the exception of the cholesterol transporter gene *NPC1*. To confirm these findings, we biotinylated the surface proteins of these gene-edited cells, immunoprecipitated with streptavidin, and performed western blotting and quantification (**Figure 4D and 4E**). A significant reduction of surface ACE2 was observed across the different cell lines except for *NPC1-edited* cells. To correlate the significance of this finding for virus infection, we edited *CCDC53*, which showed the greatest reduction in virion internalization, in Calu-3 lung cells. Viral yield was approximately ten-fold lower in the *CCDC53-*edited compared to control cells at 24 h for Sfull and 48 h for Sdel (**Figure 4F and 4G**). These results suggest retrieval and recycling complexes identified in our screen regulate expression of the ACE2 receptor, which is required for optimal SARS-CoV-2 infection.

### SARS-CoV-2 entry is elegantly regulated by endosomal cargo sorting complexes

To distinguish the complexes important for virus infection, we edited additional genes. The retriever complex is another retromer-like complex that mediates cargo recycling and consists of the genes *DSCR3, C16orf62*, and *VPS29*(McNally et al., 2017). *VPS29* and *C16orf16* that were identified in our screen, also are shared functionally by the retromer and CCC complexes(Norwood et al., 2011; Phillips-Krawczak et al., 2015).

Sorting Nexin 17 (SNX17) acts as a cargo adaptor-associated with retriever and the adaptor SNX31(McNally et al., 2017). SNX27 and SNX3 are two additional cargo adaptors associated with the retromer complex(Burd and Cullen, 2014). To test these genes, which were not identified in our screen, we introduced three sgRNAs per gene in A549-ACE2 cells and infected with Sdel virus. The editing efficiency of SNX17 and SNX27 was confirmed by western blotting (**Figure S5**). Only the retromer-associated adaptor SNX27 was required (**Figure S5**), highlighting the importance of the retromer complex over the retriever one for Sdel infection.

The COMMD proteins of CCC complex are a 10-member family (COMMD1-10)(Burstein et al., 2005) that act as cargo-binding adaptors(Bartuzi et al., 2016; Li et al., 2015). Of these 10 proteins, we identified the genes encoding all of them in our screen except for COMMD1, 6, and 9 (**Figure 2C**). Knokout of the COMMD1, 6, and 9 increases the low-density lipoprotein cholesterol levels in the plasma membrane, thereby maintaining lipid raft composition(Fedoseienko et al., 2018). In our experiments, editing each of these three genes as well as cholesterol uptake-related genes did not impact Sdel infection in A549-ACE2 or HeLa-ACE2 cells (**Figure S6A and 7B**), suggesting that these members of the COMMD protein family function differently. Notably, knockout of *COMMD1* did not affect expression of *COMMD3* or *CCDC22* in our study as opposed to previous work (**Figure S6C**)(Bartuzi et al., 2016; Fedoseienko et al., 2018). Overall, our experiments demonstrate that SARS-CoV-2 entry is regulated by endosomal cargo sorting complexes. Understanding how these complexes regulate the sorting of incoming virions might enable development of host-directed antiviral agents to control COVID-19.

### The S1/S2 boundary of spike protein impacts infection and disease in hamsters

In cell culture, we demonstrated that the Sdel virus resulted in a switch from the plasma membrane to endosomal fusion pathway for entry. In Calu-3 lung cells, which model more physiologically relevant airway epithelial cells, this switch led to a less efficient endosomal entry process. Since virus entry is the first step in establishing infection, we hypothesized that deletion at the S1/S2 boundary might reduce virus infectivity and transmissibility *in vivo*. Indeed, using the golden Syrian hamster model, a previous study showed that a SARS-CoV-2 variant with a 30-nucleotide deletion at the S1/S2 junction caused milder disease and less viral infection in the trachea and lungs compared to a virus lacking the deletion(Lau et al., 2020).

We evaluated the tissue tropism of the Sfull and Sdel virus following intranasal inoculation of golden Syrian hamsters. Nasal turbinates, trachea, lungs, heart, kidney, spleen, duodenum, brain, serum, and feces were collected. Sfull virus replicated robustly and reached peak titer at day 1 post infection, with a mean titer 31-, 126-, and 1259-fold higher than Sdel in the turbinates, trachea, and lungs, respectively (**Figure 5A**). While Sdel virus replication was delayed, no significant differences were observed by day 4 in these three tissues (**Figure 5B**). At days 2 and 4, five pieces of fresh feces were collected from each hamster. Although no infectious virus was detected by focus-forming assay (data not shown), viral RNA levels were higher in fecal samples for Sfull (20 and 40-fold) than Sdel at days 2 and 4, respectively (**Figure 5B**). Likely related to this, no infectious virus was detected in the duodenum, and Sfull RNA was 6.3-fold higher than Sdel at day 4 (**Figure S7A**). In serum, we detected no difference in viremia at day 1, but Sfull RNA was 63- and 32-fold higher than Sdel at days 2 and 4, respectively (**Figure S7B**). In other extrapulmonary organs, infectious virus was not consistently detected (data not shown). In general, brain tissue had the highest viral RNA copy number, and all organs showed higher levels of Sfull RNA at day 2 or 4 compared to Sdel except for the liver and kidneys (**Figure S7C-G**). Weight loss was only observed in hamsters inoculated with Sfull and decreased as much as ~18% at days 5 and 6 (**Figure S7H**).

**Figure 5.**
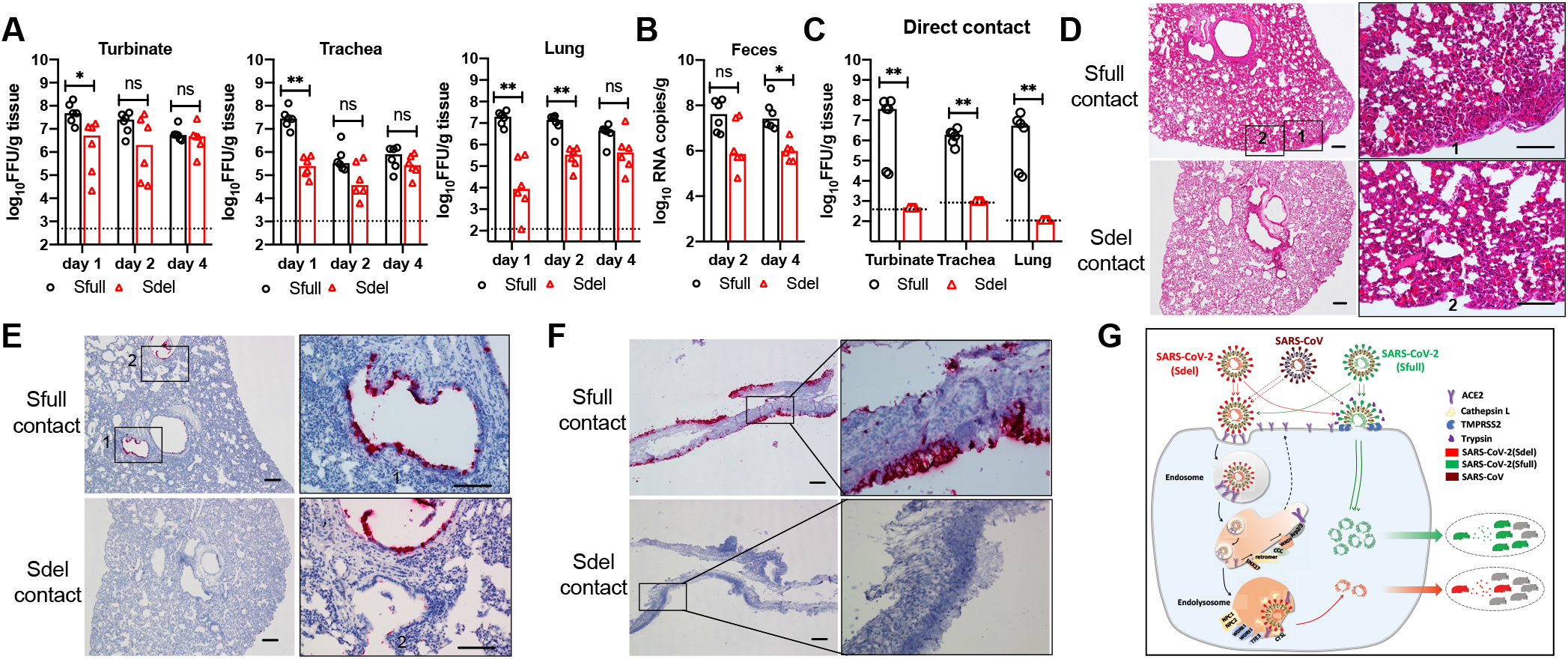
The S1/S2 boundary of SARS-CoV-2 spike protein modulates the infection and transmission in golden Syrian hamster model. **A.** Viral load in the tissues of nasal turbinate, trachea, and lung. Tissues were harvested at day 1, 2 and 4 post-challenge of Sfull or Sdel virus (n=6 per day). **B.** Viral RNA in fecal samples. Fresh fecal samples were collected at day 2 and 4 post-infection of Sfull or Sdel strain (n=6 per day) for qRT-PCR. **C.** Transmission of Sfull or Sdel strain in hamsters by direct contact exposure. Naïve hamsters (n=6) were each co-housed with one inoculated donor at day 1 for three days. Hamsters were sacrificed and the indicated tissues were harvested for titration. The dashed lines represent the limit of detection by focus-forming assay. Mean fecal sample weight (B): two-tailed unpaired t-test; median viral titers (A, B, C): two-tailed Mann–Whitney test *P < 0.05; **P < 0.01; ns, not significant. **D.** H&E staining of lung sections of contact hamsters. Representative images are shown from n = 6 hamsters. Scale bar, 100μm. **E-F.** RNA ISH of lung and nasal turbinate sections of contact hamsters. Representative images are shown from n = 6 hamsters. Scale bar, 100μm. **G.** Model of the role of S1/S2 boundary in cell entry, pathogenicity, and transmissibility of SARS-CoV-2. SARS-CoV-2 with intact spike protein (Sfull virus) preferentially enters cells at the plasma membrane (early entry pathway) in respiratory tract tissues expressing the proteases (*e.g*., TMPRSS2) to activate the membrane fusion. The virus (Sdel) with deletion at S1/S2 junction site in spike, however, tends to enter via endosomal pathway (late entry pathway). Both entry pathways are initiated with virion binding to cellular receptor ACE2 that is regulated by host factors including retromer, CCC, and WASH complexes, etc. The more efficient early entry pathway in respiratory tract with intact spike protein than the late pathway promotes virus production, pathogenesis, and transmission in a hamster model. The SARS-CoV with spike lacking the insertion of multi-basic amino acids may resemble the Sdel virus and enter cell less efficiently than SARS-CoV-2 resulting in relatively low transmissibility.

### The S1/S2 boundary of spike protein modulates the transmission

To determine the impact of deletion at the S1/S2 junction on transmissibility by direct contact exposure, six hamsters were inoculated intranasally with Sfull or Sdel virus. At 24 h post inoculation, each donor hamster was transferred to a new cage and co-housed with one naïve hamster for 3 days. For donors (day 4 post-inoculation), tissue samples were processed (**Figure 5A and 5B, and Figure S7**). For contact hamsters (day 3 post-exposure), nasal turbinate, trachea, and lungs were collected for infectious virus titration and histopathological examination. The peak titers in turbinate, trachea, and lungs from Sfull-exposed hamsters reached 8, 6.6, and 7.4 logs, respectively (6.6 logs, 6.2 logs, and 6.1 logs on average, respectively) (**Figure 5C**). Unexpectedly, no infectious virus was detected in these three tissues from Sdel-exposed hamsters (**Figure 5C**). In lung sections from hamsters that were exposed to Sfull-infected animals, we observed mononuclear cell infiltrate, protein-rich fluid exudate, hyaline membrane formation, and haemorrhage (**Figure 5D**). In contrast, no or minimal histopathological change was observed in the lung sections from hamsters that were exposed to Sdel-infected animals (**Figure 5D**). To examine viral spread in the lungs, we performed RNA *in situ* hybridization (ISH). Viral RNA was clearly detected in bronchiolar epithelial cells in hamsters exposed to Sfull-infected animals (**Figure 5E**) whereas it was rarely detected in hamsters exposed to Sdel-infected animals. Similaly, abundant RNA was observed in the nasal turbinate epithelium (**Figure 5F**). These results indicated that transmission of Sfull from infected hamsters to co-housed naïve hamsters was efficient whereas the deletion at the S1/S2 boundary in the S protein of Sdel markedly reduced transmission.

## DISCUSSION

Using authentic infectious viruses, our *in vitro* and *in vivo* studies establish that the unique S1/S2 boundary of the SARS-CoV-2 S protein can determine the entry pathways and transmission of the virus (**Figure 5G**). The Sfull virus with an intact boundary bearing the multi-basic residues, RRAR, preferentially enters cells through the plasma membrane fusion pathway, whereas Sdel with the deletion disrupting these residues switches the cell entry to a less efficient endosomal pathway. This is further demonstrated when we mutated two basic residues in the RRAR motif (R682S, R685S), which led to less efficient infection of Sdel in Calu-3 cells. In Vero cells expressing no or minimal TMPRSS2, Sfull virus enters via endosomal pathway, making the multi-basic dispensable, which results in its deletion, presumably due to an adaptive advantage. This deletion effect could be abrogated by adding trypsin or by expressing TMPRSS2, which allows the virus to resume entry via the plasma membrane fusion pathway, as we verified by the acquisition of sensitivity to camostat. In contrast, the Sdel virus maintains its usage of the E-64d-sensitive endosomal pathway for entry even in Vero-TMPRSS2 cells. The results of our experiments using the SARS-CoV spike protein, which lacks the multiple basic residues at the S1/S2 junction, were similar to what we observed for the Sdel virus. It is noteworthy that infection by Sdel virus, but not Sfull, in A549-ACE2 cells is sensitive to the cathepsins B/L inhibitor E-64d, highlighting the importance of S1/S2 boundary sequence in this entry process. Treatment with camostat has no impact on Sfull virus infection in A549-ACE2 cells, as no or minimal TMPRSS2 is expressed, suggesting that other TMPRSS2 homologs or trypsin-like proteases may activate the Sfull virus entry at the plasma membrane.

The notion that this deletion at the S1/S2 boundary discriminates the entry pathway used by the virus was supported by the large number of endosomal entry host factors uncovered in our genome-wide CRISPR screen. Genes for the endosomal entry-specific enzyme CTSL and for regulating endolysomal trafficking and membrane fusion, such as *NPC1/2* and *WDR81/91*, were required for Sdel, but not for Sfull virus infection. In parallel, we discovered a panel of entry factors common to both Sdel and Sfull that regulate the surface expression of the SARS-CoV-2 receptor ACE2. Understanding the detailed mechanisms of action for these common host factors could help in the development of potential countermeasures to combat COVID-19. More importantly, because Sfull virus preferentially enters cells at the plasma membrane, targeting the endosomal entry pathway might not be a promising strategy to inhibit SARS-CoV-2 infection. This is exemplified by the *in vitro* and *in vivo* results of studies examining the lysosomal acidification inhibitors chloroquine and hydroxychloroquine(Boulware et al., 2020; Hoffmann et al., 2020c; Kupferschmidt).

The serine protease TMPRSS2 on the cell surface activates the spike protein-mediated membrane fusion pathway, which is important for virus spread(Iwata-Yoshikawa et al., 2019; Zhou et al., 2015). It has been reported that TMPRSS2 is enriched in nasal and bronchial tissues(Qi et al., 2020; Sungnak et al., 2020a; Sungnak et al., 2020b), implying that the transmission of SARS-CoV-2 by respiratory droplets might be enhanced for virus bearing an intact versus a deleted S1/S2 boundary. In our hamster experiments, the deletion mutant virus Sdel exhibited decreased viral infection and disease compared to Sfull. More importantly, the transmission of Sdel by direct contact exposure for 3 days was almost completely abrogated. The nearly complete abrogation of infection by direct contact highlights the critical role of the multi-basic sequence at the S1/S2 boundary in transmissibility, presumably due to usage of the more efficient fusion entry pathway.

## METHODS

### Cells

Vero E6 (Cell Bank of the Chinese Academy of Sciences, Shanghai, China), HEK 293T (ATCC # CRL-3216), HeLa (ATCC #CCL-2), A549 (kindly provided by M.S. Diamond, Washington University), and Calu-3 (Cell Bank of the Chinese Academy of Sciences, Shanghai, China) all were cultured at 37°C in Dulbecco’s Modified Eagle Medium (Hyclone #SH30243.01) supplemented with 10% fetal bovine serum (FBS), 10 mM HEPES, 1 mM Sodium pyruvate, 1× non-essential amino acids, and 100 U/ml of Penicillin-Streptomycin. The A549-ACE2 and HeLa-ACE2 clonal cell lines were generated by transduction of lentivector expressing the human ACE2 gene as described bellow. Similarly, the bulk Vero-TMPRSS2 cells were generated by transduction of lentivector expressing the human TMPRSS2 and selected with puromycin. The surface expression of ACE2 or TMPRSS2 was confirmed by flow cytometry. All cell lines were tested routinely and free of mycoplasma contamination.

### Viruses

The SARS-CoV-2 nCoV-SH01 strain (GenBank accession no. MT121215) was isolated from a COVID-19 patient by passaging in Vero E6 cells twice in the presence of trypsin. This virus stock underwent three rounds of plaque-purification in Vero E6 cells in the presence of trypsin and designated as SH01-Sfull (thereafter as Sfull). Sfull stain was then passaged twice and plaque-purified once in the absence of trypsin, resulting the stain Sdel that has 21 nt deletion in the spike gene. Sfull virus was also passaged twice in Vero E6 cells in the presence of trypsin or twice in Vero E6 ectopically expressing the TMPRSS2 without trypsin. The virus titers were titrated in Vero E6 cells in the presence of trypsin by focus-forming assay as described below. The full-genome of Sfull and Sdel strains, and the entire spike gene of other passaged viral stocks were Sanger sequenced and analyzed. All the sequencing primers are available upon request. All experiments involving virus infections were performed in the biosafety level 3 (BSL-3) facility of Fudan University following the regulations.

### Genome-wide CRISPR sgRNA screen

A human Brunello CRISPR knockout pooled library encompassing 76,441 different sgRNAs targeting 19,114 genes(Doench et al., 2016) was a gift from David Root and John Doench (Addgene #73178), and amplified in Endura cells (Lucigen #60242) as described previously(Joung et al., 2017; Sanjana et al., 2014). The sgRNA plasmid library was packaged in 293FT cells after co-transfection with psPAX2 (Addgene #12260) and pMD2.G (Addgene #12259) at a ratio of 2:2:1 using Fugene^®^HD (Promega). At 48 h post transfection, supernatants were harvested, clarified by spinning at 3,000 rpm for 15 min, and aliquotted for storage at - 80°C.

For the CRISPR sgRNA screen, A549-ACE2-Cas9 cells were generated by transduction of A549-ACE2 cell line with a packaged lentivirus expressing the mCherry derived from the lentiCas9-Blast (Addgene #52962) that the blasticidin resistance gene was replaced by mCherry. The sorted mCherry positive A549-ACE2-Cas9 cells were transduced with packaged sgRNA lentivirus library at a multiplicity of infection (MOI) of ~0.3 by spinoculation at 1000g and 32°C for 30 min in 12-well plates. After selection with puromycin for around 7 days, ~1 x 10^8^ cells in T175 flasks were inoculated with SARS-CoV-2 Sdel strain (MOI of 3) and then incubated until nearly all cells were killed. The medium was changed and remaining live cells grew to form colonies. The cells were then harvested and re-plated to the flasks. After second round of killing by the virus, the remaining cells were expanded and ~3×10^7^ of cells were collected for genomic DNA extraction. Genomic DNA from the uninfected cells (5 × 10^7^) was extracted as the control. The sgRNA sequences were amplified(Shalem et al., 2014) and subjected to next generation sequencing using an Illumina NovaSeq 6000 platform. The sgRNA sequences targeting specific genes were extracted using the FASTX-Toolkit (http://hannonlab.cshl.edu/fastx_toolkit/) and cutadapt 1.8.1, and further analyzed for sgRNA abundance and gene ranking by a published computational tool (MAGeCK)(Li et al., 2014) (see Supplementary Tables 1 and 2).

### Gene validation

Top 35 genes from the MAGeCK analysis were selected for validation. Two independent sgRNAs per gene were chosen from the Brunello CRISPR knockout library and cloned into the plasmid lentiCRISPR v2 (Addgene #52961) and packaged with plasmids psPAX2 and pMD2.G. A549-ACE2, HeLa-ACE2, or Calu-3 cells were transduced with lentiviruses expressing individual sgRNA and selected with puromycin for 7 days. The gene-edited mixed population of cells was used for all the experiments in this study.

For virus infection, gene-edited A549-ACE2 or HeLa-ACE2 cells were inoculated with Sfull (MOI 2) and Sdel (MOI 2). Vero, Vero-TMPRSS2, and Calu-3 cells were inoculated with Sfull (MOI 1) and Sdel (MOI 1). At 24 h post infection, cells were fixed with 4% paraformaldehyde (PFA) diluted in PBS for 30 min at room temperature, and permeabilized with 0.2% Triton x-100 in PBS for 1 h at room temperature. Cells then were subjected for immunofluorescence staining and imaging as described bellow. Validation also was performed by an infectious virus yield assay.

### Virus yield assay

Vero, Vero-TMPRSS2, A549-ACE2, and Calu-3 cells were seeded one day prior to infection. Cells were inoculated with same MOI of Sfull or Sdel (Vero, Vero-TMPRSS2, MOI 0.01; A549-ACE2, MOI 2; Calu-3, MOI 0.1) for 1 h. After three times of washing, cells were maintained in 2% FBS culture media, and supernatants were collected at specific time points for titration on Vero cells by focus-forming assay.

### Pseudotyped virus experiment

Pseudoviruses were packaged in HEK 293T cells by co-transfecting the retrovector pMIG (kindly provided by Jianhua Li, Fudan Univiersity) for which the gene of target was replaced by the nanoluciferase gene, plasmid expressing the MLV Gag-Pol, and pcDNA3.1 expressing different spike genes or VSV-G (pMD2.G (Addgene #12259)) using Fugene^®^HD tranfection reagent (Promega). At 48 h post transfection, the supernatant was harvested, clarified by spinning at 3500 rpm for 15 min, aliquoted and stored at −80C for use. The virus entry was assessed by transduction of pseudoviruses in gene-edited cells in 96-well plates. After 48 or 72 h, the luciferase activity was determined using Nano-Glo^®^ Luciferase Assay kit (Promega #N1110) according to the manufacturer’s instructions. The same volume of assay reagent was added to each well and shake for 2 min, After incubation at room temperature for 10 min, luminescence was recorded by using a FlexStation 3 (Molecular Devices) with an integration time of 1 second per well.

### Plasmid construction

To construct the lentivector expressing the human ACE2 gene, the human ACE2 gene (Miaolingbio #P5271) was PCR-amplified and cloned into the pLV-EF1a-IRES-blast (Addgene #85133). The human TMPRSS2 and DPP4 gene (Sino Biological #HG13070-CM) was cloned by the similar strategy. To construct the vectors for pseudovirus packaging, the full-length spike gene was PCR-amplified from Sfull or Sdel strain and cloned into the pcDNA3.1 vector. The Sfull spike gene with two mutations (R682S, R685S)(Wang et al., 2020a) in the furin cleavage site was generated by PCR. The full-length SARS-CoV or MERS-CoV spike gene was cloned similarly.

### Virus binding and internalization assays

A549-ACE2 gene-edited cells were seeded in 24-well plate one day prior to the assays. Plates were pre-incubated on ice for 10 min, then washed twice with ice-cold PBS. Ice-cold Sfull virus (MOI of 5) in a 0.5-ml medium was incubated with cells on ice for 45 min. After five cycles of washing, cells were lysed in TRIzol reagent (ThermoFisher #15596018) for RNA extraction. For internalization assay, after 5 cycles of washing, cells were incubated into medium supplemented with 2% FBS and then incubated at 37°C for 45 min. Cells were chilled on ice, washed with ice-cold PBS, and then treated with 400 μg/ml protease K on ice for 45 min. After three additional washes, cells were lysed in TRIzol reagent for RNA extraction. RT-qPCR was conducted to quantify the viral specific nucleocapsid RNA and an internal control GAPDH.

### Cell-based S1-Fc and anti-ACE2 antibody binding assay

A549-ACE2 gene-edited cells were seeded in 96-well plate one day prior to the experiment. Cells were collected with TrypLE (Thermo #12605010) and washed twice with ice-cold PBS. Live cells were incubated with the recombinant protein, S1 domain of SARS-CoV-2 spike C-terminally fused with Fc (Sino Biological #40591-V02H, 1μg/ml), or the anti-ACE2 antibody (Sino Biological #10108-RP01, 1 μg/ml) at 4 °C for 30 min. After washing, cells were stained with goat anti-human IgG (H + L) conjugated with Alexa Fluor 647 (Thermo #A21445, 2 μg/ml) for 30 min at 4 °C. After two additional washes, cells were subjected to flow cytometry analysis (Thermo, Attune^™^ NxT).

### Western blotting

Cells in plates washed twice with ice-cold PBS and lysed in RIPA buffer (Cell Signaling #9806S) with a cocktail of protease inhibitors (Sigma-Aldrich # S8830). Samples were prepared in reducing buffer (50 mM Tris, pH 6.8, 10% glycerol, 2% SDS, 0.02% [wt/vol] bromophenol blue, 100 mM DTT). After heating (95°C, 10 min), samples were electrophoresed in 10% SDS polyacrylamide gels, and proteins were transferred to PVDF membranes. Membranes were blocked with 5% non-fat dry powdered milk in TBST (100mM NaCl, 10mM Tris,pH7.6, 0.1% Tween 20) for 1 h at room temperature, and probed with the primary antibodies at 4 C overnight. After washing with TBST, blots were incubated with horseradish peroxidase (HRP)-conjugated secondary antibodies for 1 h at room temperature, washed again with TBST, and developed using SuperSignal West Pico or Femto chemiluminescent substrate according to the manufacturer’s instructions (ThermoFisher). The antibodies used are as follows: rabbit anti-COMMD3 (proteintech #26240-1-AP, 1:800), rabbit anti-VPS35 (proteintech #10236-1-AP,1:500), rabbit anti-CCDC22 (proteintech #16636-1-AP, 1:1000), rabbit anti-NPC1 (proteintech #13926-1-AP, 1:1000), rabbit anti-NPC2 (proteintech #19888-1-AP, 1:800), rabbit anti-CCDC53 (proteintech #24445-1-AP, 1:500), rabbit anti-COMMD1 (proteintech #11938-1-AP, 1:2000), mouse anti-SNX27 (Abcam #ab77799, 1:1000), rabbit anti-SNX17 (proteintech, #10275-1-AP, 1:2000), rabbit anti-LDLR (proteintech, #10785-1-AP, 1:1000), rabbit anti-LRP1 (Abcam #ab92544, 1:5000), rabbit anti-SARS-Cov-2 spike S2 (Sino Biological #40590-T62, 1:1000), rabbit anti-β-actin (proteintech #20536-1-AP, 1:2000). The HRP-conjugated secondary antibodies include: Goat anti-mouse (sigma #A4416, 1:5000), goat anti-rabbit (thermo fisher #31460, 1:5000), goat anti-human (sigma #A6029, 1:5000).

For quantification studies, after probing with primary antibodies, membranes were incubated with goat anti-rabbit IRDye 800CW secondary antibody (LI-COR #926-32211, 1:10000), goat anti-rabbit IRDye 680RD secondary antibody (LI-COR #926-68071, 1:10000) or goat anti-mouse IRDye 800CW secondary antibody (LI-COR #926-32210, 1:10000), then developed and analyzed with the Odyssey CLx Imaging System.

### Biotinylation of plasma membrane proteins

Gene-edited A549-ACE or Calu-3 cells seeded in 6-well plate 24 h prior to experiment were chilled on ice for 10 min, and labeled with 2.5 mg/ml Biotin (Thermo fisher #21331) in PBS for 30 min on ice. Cells were quenched with 100 mM glycine in PBS 3 times, 10 min each. After washing with PBS, cells were lysed in RIPA buffer (Cell Signaling #9806S) with a cocktail of protease inhibitors (Sigma-Aldrich # S8830), and immunoprecipitated with Streptavidin agarose beads overnight at 4°C. Beads were then washed three times with RIPA buffer, and eluted into 5x loading buffer (Beyotime #P0015L) at 95°C for 10min. After spinning at maximum speed for 10 min, the supernatants were harvested for western blotting using rabbit anti-ACE2 (Abcam #ab15348, 1:1000) as described above, and analyzed with the Odyssey CLx Imaging System. The un-immnoprecipitated lysates were used as loading control.

### Immunofluorescence assay

Virus-infected cells were washed twice with PBS, fixed with 4% paraformaldehyde in PBS for 30 min, permeablized with 0.2% Triton X-100 for 1 h. Cells were then incubated with house-made mouse anti-SARS-CoV-2 nucleocapsid protein serum (1:1000) at 4 °C overnight. After three washes, cells were incubated with the secondary goat anti-mouse antibody conjugated with Alexa Fluor 555 (Thermo #A-21424, 2 μg/ml) for 2 h at room temperature, followed by staining with 4’,6-diamidino-2-phenylindole (DAPI). Images were collected using an Operetta High Content Imaging System (PerkinElmer), and processed using the ImageJ program (http://rsb.info.nih.gov/ij/).

### Cell viability assay

A CellTiter-Glo^®^ Luminescent Cell Viability Assay (Promega # G7570) was performed according to the manufacturer’s instructions. The same number of gene-edited cells was seeded into opaque-walled 96-well plates. 48 h later, CellTiter-Glo^®^ reagent was added to each well and allowed to shake for 2 min. After incubation at room temperature for 10 min, luminescence was recorded by using a FlexStation 3 (Molecular Devices) with an integration time of 0.5 second per well.

### Animal experiments

Six to ten week-old male hamsters were used in the study in the BSL-3 laboratory of Fudan University. The experiment protocol has been approved by the Animal Ethics Committee of School of Basic Medical Sciences at Fudan University. The hamsters were inoculated intranasally with 5×10^4^ focus-forming unit (FFU) of Sfull or Sdel virus. To evaluate the viral transmission by direct contact, at day 1 post-infection, each hamster infected with Sfull or Sdel was transferred to a new cage and co-housed with one age-matched naïve hamster for three days. At 24 h, 48 h, and 96 h post virus challenge, or 72 h post contact, animals were euthanized and the sera were collected. After perfusion extensively with PBS, indicated tissues were harvested for virus titration by focus-forming assay in the presence of trypsin or histopathological examination. To collect fecal samples, at 48 h and 96 h post challenge, each hamster was put into an individual clean container and fresh fecal samples (5 pieces) were collected and frozen down for virus titration by focus-forming assay or RT-qPCR analysis. To monitor the body weight change, hamsters were measured daily for 14 days. Tissues were homogenized in DMEM and virus was titrated by focus-forming assay (FFA)(Pal et al., 2013) using the rabbit polyclonal antibody against SARS-CoV nucleocapsid protein (Rockland, 200-401-A50, 0.5μg/ml) or by RT-qPCR after RNA extraction as described below.

### Histology and RNA *in situ* hybridization

Virus-infected hamsters were euthanized and perfused extensively with PBS. Nasal turbinate and lung tissues were harvested and fixed in 4% paraformaldehyde (PFA) for 48 h. Tissues were embedded in paraffin for sectioning and stained with hematoxylin and eosin (H&E) to assess tissue morphology. To determine sites of virus infection, RNA *in situ* hybridization was performed using the RNAscope 2.5 HD Assay (Red Kit) according to the manufacturer’s instructions (Advanced Cell Diagnostics). In brief, sections were deparaffinized, treated with H_2_O_2_ and Protease Plus prior to probe hybridization. A probe specifically targeting the SARS-CoV-2 spike RNA (Advanced Cell Diagnostics, #848561) was used for *in situ* hybridization (ISH) experiments. Tissues were counterstained with Gill’s hematoxylin. Tissue sections were visualized using a Nikon Eclipse microscope.

### qRT-PCR

RNA from serum, tissues, or cells was extracted with the TRIzol reagent (ThermoFisher #15596018). Viral or host RNA levels were determined using the TaqPath™ 1-Step RT-qPCR Master Mix (ThermoFisher # A15299) on CFX Connect Real-Time System (Bio-Rad) instrument. A standard curve was produced using serial 10-fold dilutions of *in vitro* transcribed RNA of N gene driven by the SP6 promoter (ThermoFisher #AM1340). Viral burden was expressed on a log10 scale as viral RNA copies per g of tissue or ml of serum. Primers and probes used are as follows: nCoV-N-Fwd: 5’-GACCCCAAAATCAGCGAAAT-3’; nCoV-N-Rev: 5’-TCTGGTTACTGCCAGTTGAATCTG-3’; nCoV-N-Probe: 5’-FAM-ACCCCGCATTACGTTTGGTGGACC-BHQ1-3’; hGAPDH-Fwd: 5’-TGCCTTCTTGCCTCTTGTCT-3’; hGAPDH-Rev: 5’-GGCTCACCATGTAGCACTCA-3’; and GAPDH-Probe: 5’-FAM-TTTGGTCGTATTGGGCGCCTGG-BHQ1-3’.

### Virus load determination by focus-forming assay

The experiment was performed similarly as previously described(Brien et al., 2013). Briefly, Vero E6 monolayer in 96-well plates were inoculated with serially diluted virus for 2 h and then overlaid with methylcellulose for 48 h. Cells were fixed with 4% paraformaldehyde in PBS for 1 h and permeablized with 0.2% Triton X-100 for 1 h. Cells were stained with rabbit polyclonal antibody against SARS-CoV nucleocapsid protein (Rockland, 200-401-A50, 0.5μg/ml) overnight at 4°C, incubated with the secondary goat anti-rabbit HRP-conjugated antibody for 2 h at room temperature. The focus-forming unit was developed using TrueBlue substrate (Sera Care #5510-0030).

### Statistical analysis

Statistical significance was assigned when *P* values were < 0.05 using Prism Version 8 (GraphPad). Data analysis was determined by a Mann-Whitney, or ANOVA, or unpaired t-test depending on data distribution and the number of comparison groups.

## Data Availability

The authors declare that all data supporting the findings of this study are available within the paper and its Supplementary information. The Supplemental Tables provide data for the CRISPR-Cas9 screen, statistical analysis.

## ACKNOWLEDGEMENTS

Grants from the National Natural Science Foundation of China (32041005 to R.Z.), National Key Research and Development Program of China (2020YFA0707701 to R.Z.), Project of Novel Coronavirus Research of Fudan University (to Y.X.), and Development Programs for COVID-19 of Shanghai Science and Technology Commission (20431900401) supported this work. We thank Prof. Michael S. Diamond (Washington University) for discussions and editorial comments on the manuscript. We also thank Prof. Bin Zhou (Nanjing Agricultural University) for providing key reagents.

We wish to acknowledge colleagues at the Biosafety Level 3 Laboratory of Fudan University for help with experiment design and technical assistance.

## AUTHOR CONTRIBUTIONS

Y.Z., F.F., G.H., Y.W., Y.Y., Y.Z., W.X., R.Z. performed the experiments. Y.Z., F.F., G.H., Y.W., R.Z. designed the experiments. X.C., Z.S., W.H., Q.D., H.C., Q.C., D.Q., Y.X., Z.Y. provided administrative, supervision, technical, or material support. Y.Z., F.F., G.H., Y.W., Y.Y., R.Z. performed data analysis. R.Z. wrote the initial draft of the manuscript, with the other authors contributing to editing into the final form.

## COMPETING FINANCIAL INTERESTS

None.

## FIGURE SLEGENDS

**Figure S1.**
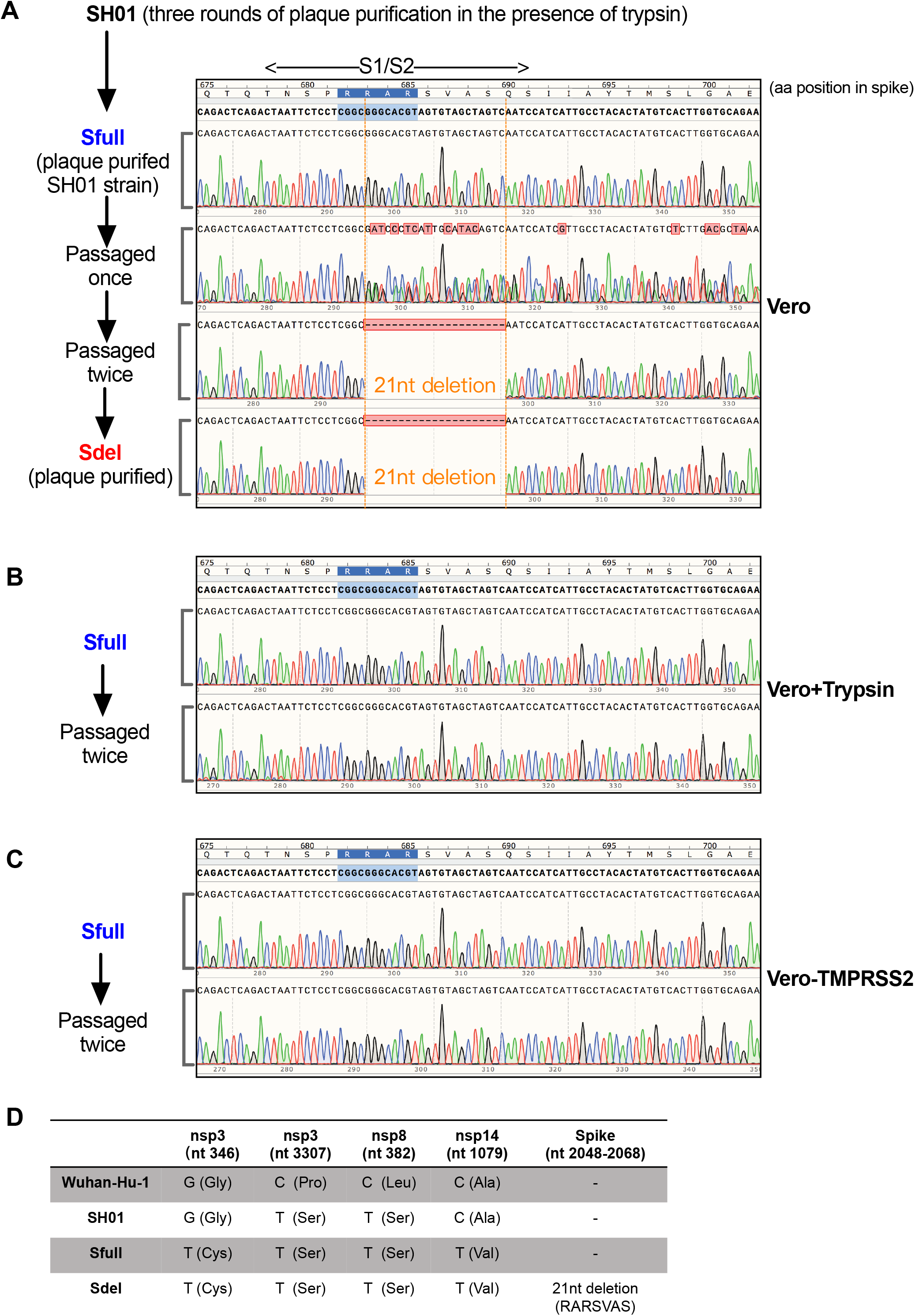
The acquisition of Sfull and Sdel clones of SARS-CoV-2. **A.** SARS-CoV-2 SH01 strain isolated from a patient sample was purified three times by plaque assay on Vero-E6 (thereafter as Vero) cells in the presence of trypsin, resulting the clone of Sfull virus. The Sdel clone was obtained by passaging the Sfull virus twice and plaque-purified once on Vero cells without trypsin. The trace results of Sanger sequencing were generated by SnapGene Viewer, and the 21 nucleotide (nt) deletion was indicated. **B.** The Sfull strain was passaged twice on Vero cells in the presence of trypsin. **C.** The Sfull strain was passaged twice on Vero cells expressing the TMPRSS2 in the absence of trypsin. d. The sequence alignment of SARS-CoV-2 strains. The full-length genome sequences obtained by RT-PCR and Sanger sequencing were aligned and compared to the stain Wuhan-Hu-1. Wuhan-Hu-1, accession No. MN908947; SH01, accession No. MT121215.

**Figure S2.**
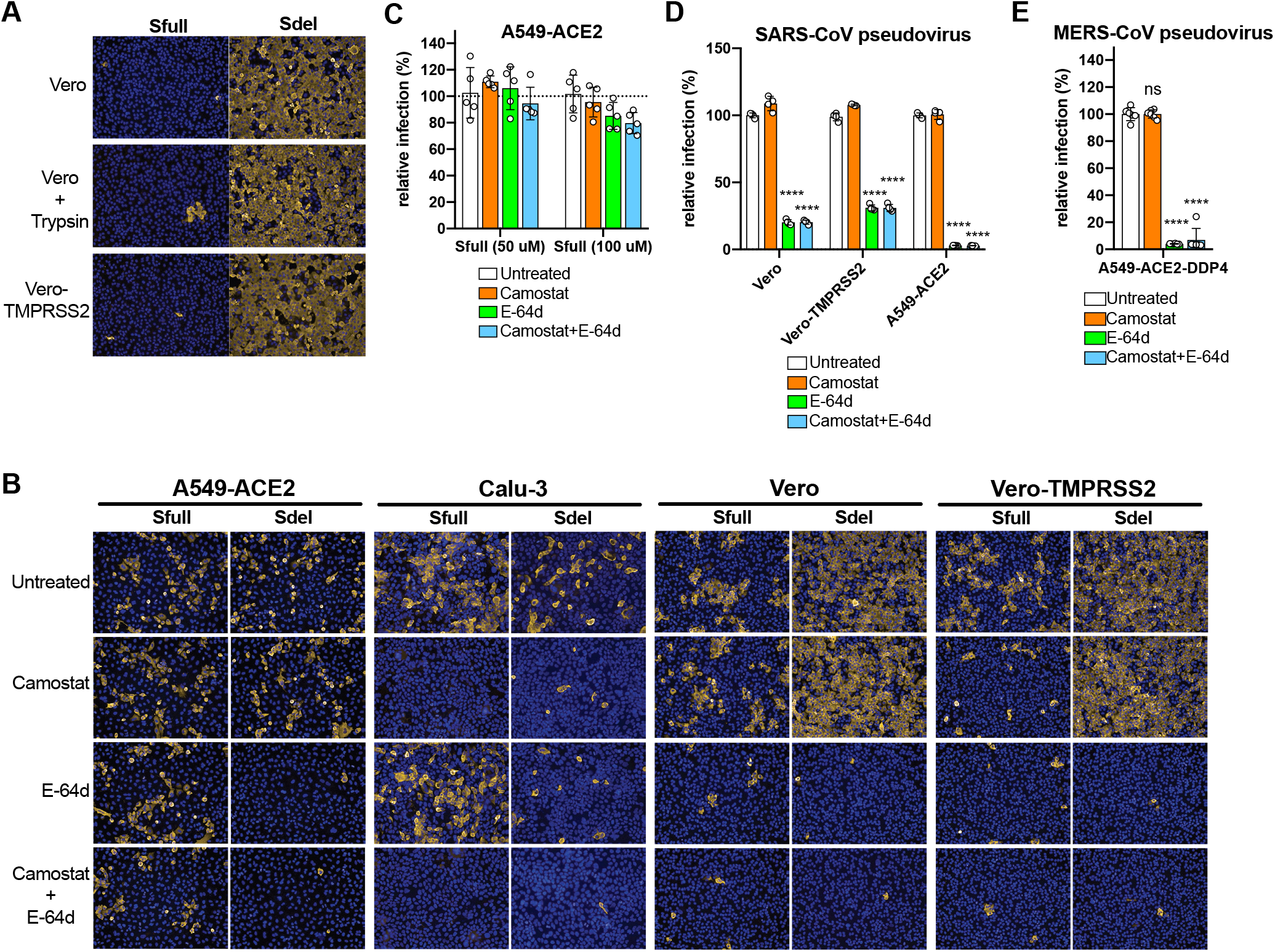
The replication and entry property of Sfull and Sdel clones of SARS-CoV-2 and SARS-CoV pseudovirus. **A.** Immunofluoresence staining of the nucleocapsid (N) protein of Sfull or Sdel virus on wild type Vero, Vero with trypsin treatment, and Vero expressing the TMPRSS2 cells. Virus-infected cells were fixed, permeablized, and stained with the house-made mouse anti-nucleocapsid serum. After washing, cells were incubated with goat anti-mouse antibody conjugated with Alexa Fluor 555 (Thermo # A-21424, 2 μg/ml), followed by staining with 4’,6-diamidino-2-phenylindole (DAPI). Images were collected using an Operetta High Content Imaging System (PerkinElmer), and processed using the ImageJ software. **B.** The effect of compounds Comostat and E-64d on the infection by Sfull or Sdel virus in different cell types. Cells were pretreated with 25 μM compounds Camostat or / and E-64d and infected with Sfull or Sdel virus in the presence of compounds for 24 h. Immunofluoresence assay was conducted as described above. **C.** Sfull infection on A549-ACE2 cells were resistant to the treatment of 50 μM or 100 μM of Comostat or / and E-64d. Cells were pretreated with the indicated compounds and infected with the Sfull virus in the presence of compounds, followed by Immunofluoresence staining as described above. **D-E.** The effect of compounds Comostat and E-64d on the infection by SARS-CoV or MERS-CoV pseudovirus in different cell types. Cells were pretreated with 25 μM compounds Camostat or / and E-64d and infected with Sfull or Sdel pseudovirus in the presence of compounds for 48 h. One-way ANOVA with Dunnett’s test *P < 0.05; ***, P < 0.001; ****P < 0.0001; ns, not significant. Immunofluoresence assay was conducted as described above. Data were pooled from two independent experiments performed in triplicate, and are normalized to the controls of individual experiments.

**Figure S3.**
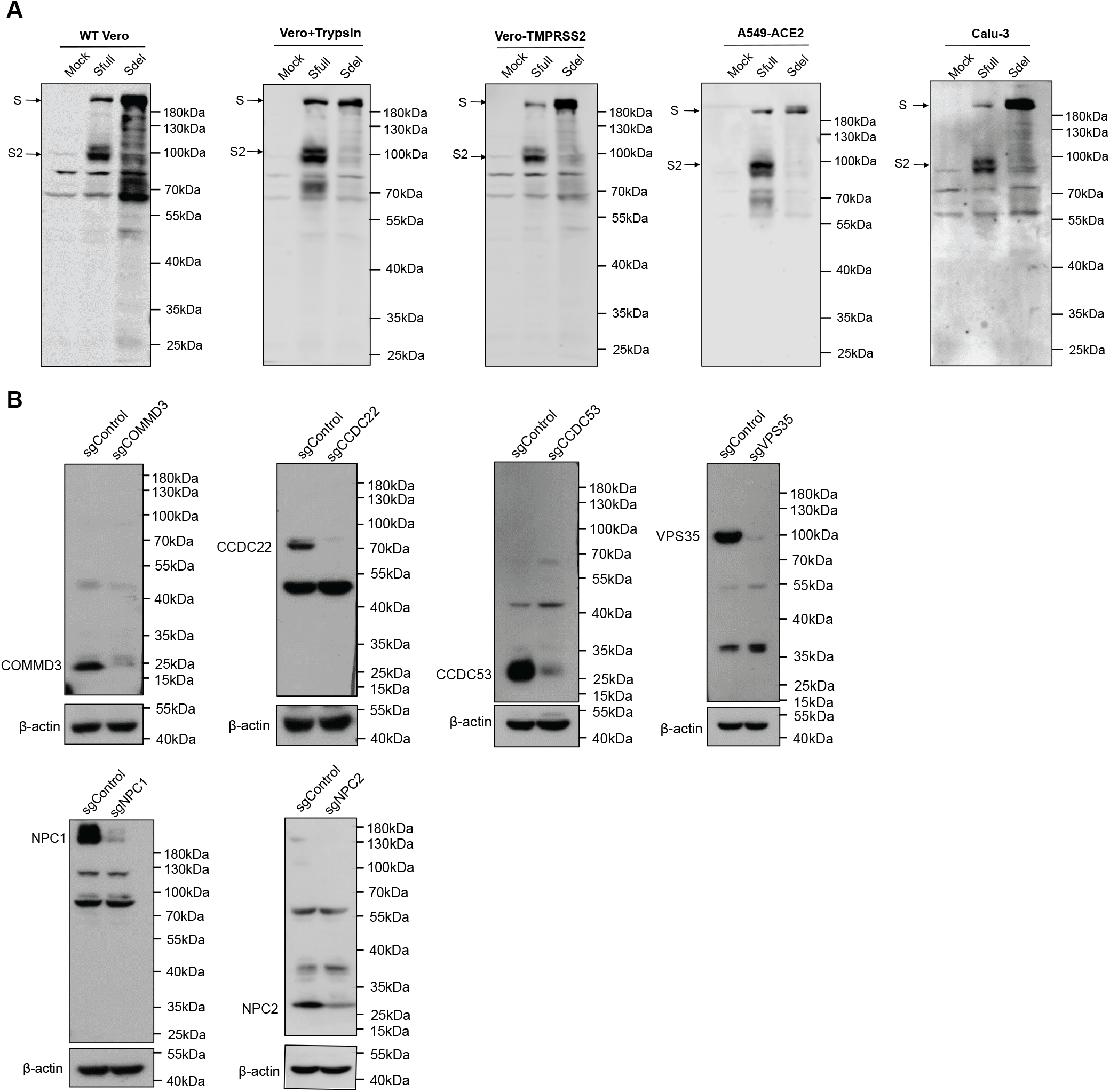
The cleavage of spike protein in different cell types and editing efficiency of A549-ACE2 cells by CRISPR sgRNA of genes selected. **A**. Western blotting of cell lysates of different cell types or conditions inoculated with Sfull or Sdel virus. The cell lysates were probed with rabbit anti-SARS-CoV-2 spike S2 anbitody (Sino Biological #40590-T62), followed by incubating with horseradish peroxidase (HRP)-conjugated goat anti-rabbit polyclonal antibody and developed using SuperSignal West Pico chemiluminescent substrate. The bands corresponding to the full-length spike (S) and cleaved S2 subunit are indicated by arrows**. B.** Editing efficiency of A549-ACE2 cells by CRISPR sgRNA of genes selected. Genes in A549-ACE2 cells were edited by the indicated sgRNAs and the mixed population of cells was subjected to western blotting. Cell lysates were probed with rabbit anti-COMMD3 (proteintech #26240-1-AP), CCDC22 (proteintech #16636-1-AP), CCDC53 (proteintech #24445-1-AP), VPS35 (proteintech #10236-1-AP), NPC1 (proteintech #13926-1-AP), or NPC2 (proteintech #19888-1-AP) polyclonal antibody, followed by incubating with horseradish peroxidase (HRP)-conjugated goat anti-rabbit polyclonal antibody and developed using SuperSignal West Pico or Femto chemiluminescent substrate.

**Figure S4.**
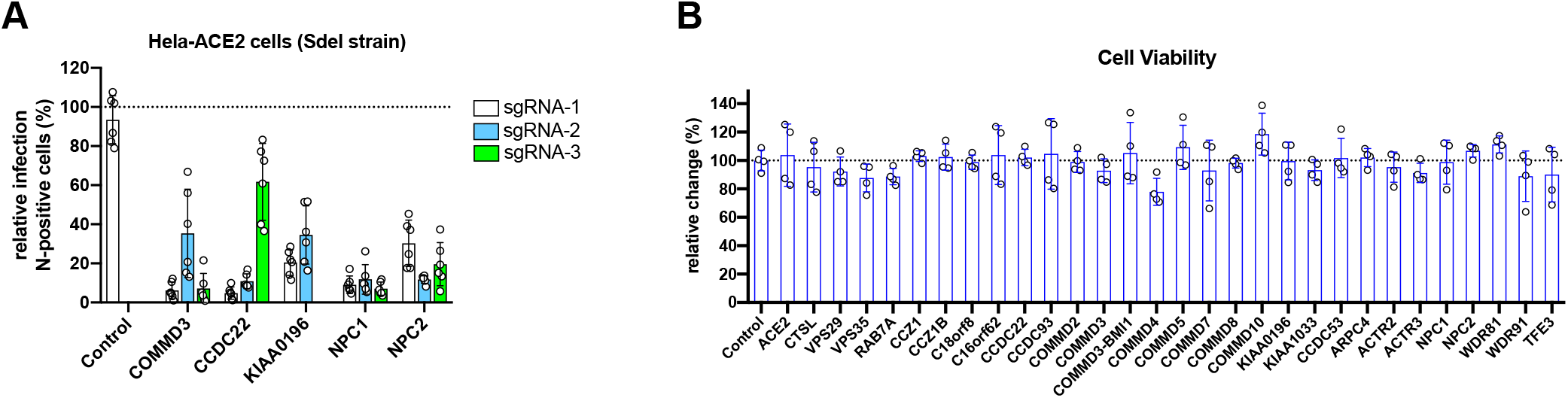
Validation of virus infection and cell viability in HeLa-ACE2 or A549-ACE2 cells. **A.** HeLa cells expressing the human ACE2 were edited by three different sgRNAs of selected genes. Cells were infected with Sdel virus and subjected to immunofluorescence assay and high-content imaging as described in Methods. Data were pooled from two independent experiments performed in triplicate, and are normalized to the controls of individual experiments. **B.** Viability of A549-ACE2 cells edited with individual CRISPR sgRNA of genes selected. An equal number of cells were plated and viability was assessed over a 48 h period using the Cell-Titer Glo assay. The results were normalized to control cells and are pooled from two independent experiments performed in duplicate.

**Figure S5.**
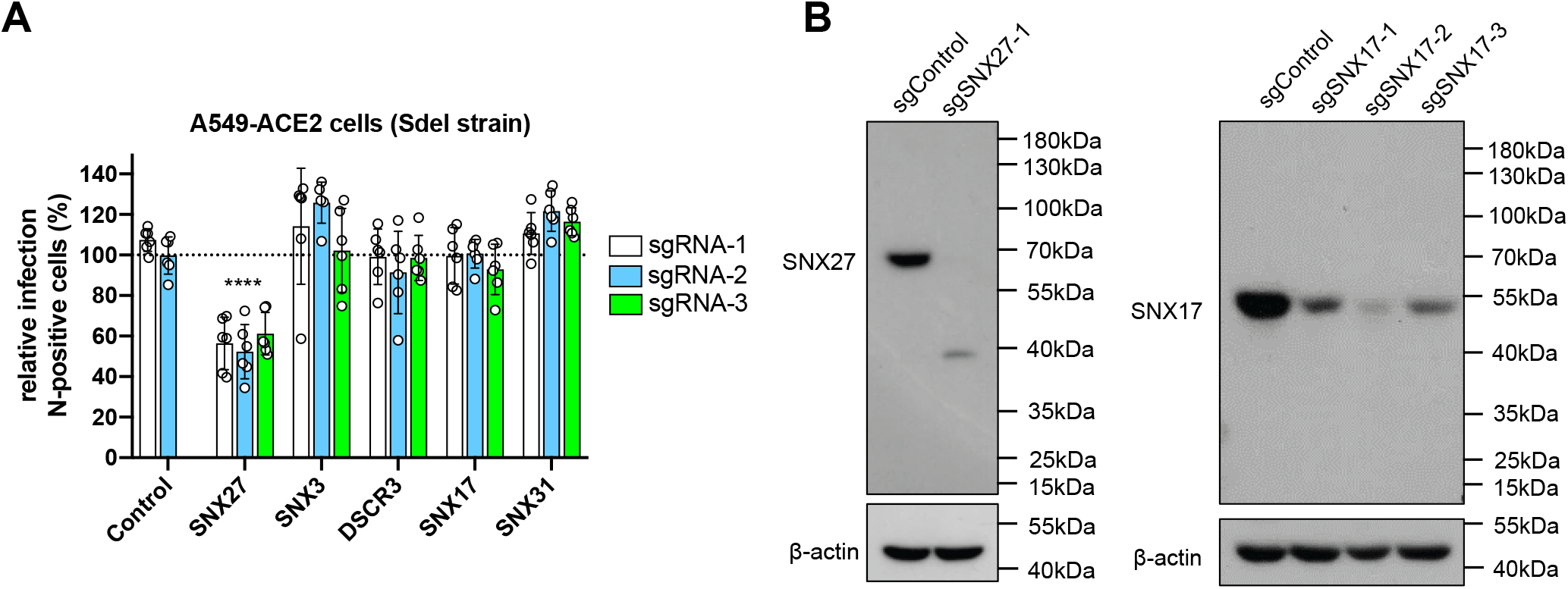
Validation of genes related to cargo retrieval and recycling in A549-ACE2 cells. **A.** A549 cells expressing the human ACE2 were edited by three different sgRNAs of selected genes. Cells were infected with Sdel virus and subjected to immunofluorescence assay and high-content imaging as described in Methods. Data were pooled from two independent experiments performed in triplicate, and are normalized to the controls of individual experiments. One-way ANOVA with Dunnett’s test ****P < 0.0001, mean ± SD. **B.** Western blotting to confirm the editing efficiency in A549-ACE2 cells by the indicated sgRNAs. Cell lysates were probed with mouse anti-SNX27 (Abcam #ab77799) or rabbit anti-SNX17 (proteintech, #10275-1-AP) polyclonal antibody, followed by incubating with horseradish peroxidase (HRP)-conjugated goat anti-mouse or rabbit polyclonal antibody, and developed using SuperSignal West Pico or Femto chemiluminescent substrate.

**Figure S6.**
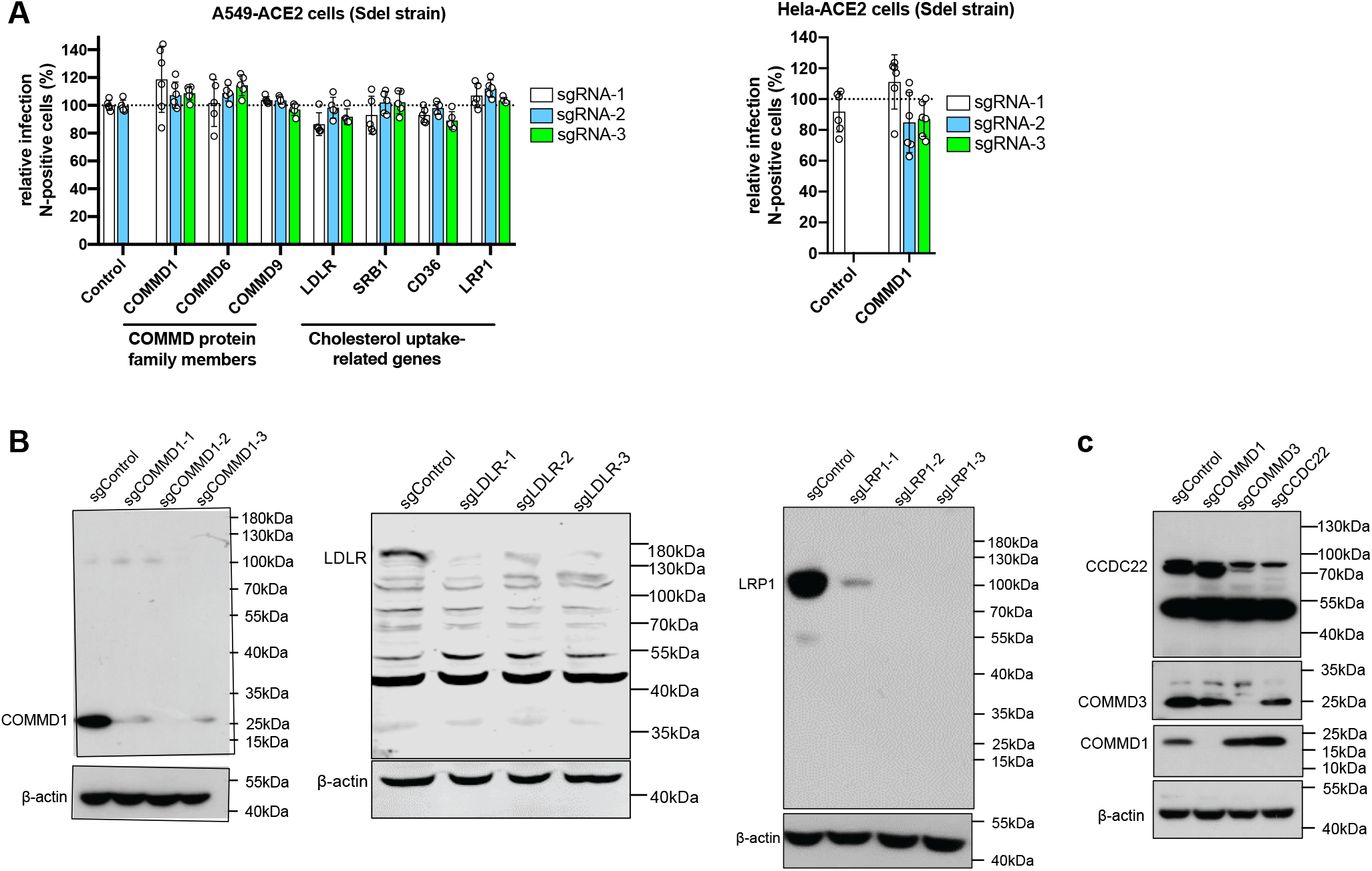
Validation of COMMD protein family members and cholesterol uptake related genes in A549-ACE2 or HeLa-ACE2 cells. **A.** COMMD1, 6, and 9, and genes related to cholesterol uptake are not required for Sdel virus infection. A549 or HeLa cells expressing the human ACE2 were edited by three different sgRNAs of the indicated genes. Cells were infected with Sdel virus and subjected to immunofluorescence assay and high-content imaging as described in Methods. Data were pooled from two independent experiments performed in triplicate, and are normalized to the controls of individual experiments. **B.** Western blotting to verify the editing efficiency of COMMD1, LDLR, and LRP1 genes in A549-ACE2 cells by the indicated sgRNAs. **c.** Western blotting to verify the expression of COMMD1, COMMD3, or CCDC22 affected by gene-editing. Cell lysated were probed with rabbit anti-COMMD1 (proteintech #11938-1-AP), LDLR (proteintech, #10785-1-AP), LRP1 (Abcam #ab92544), rabbit anti-COMMD3 (proteintech #26240-1-AP), or rabbit anti-CCDC22 (proteintech #16636-1-AP) polyclonal antibody, followed by incubating with horseradish peroxidase (HRP)-conjugated goat anti-rabbit polyclonal antibody and developed using SuperSignal West Pico or Femto chemiluminescent substrate.

**Figure S7.**
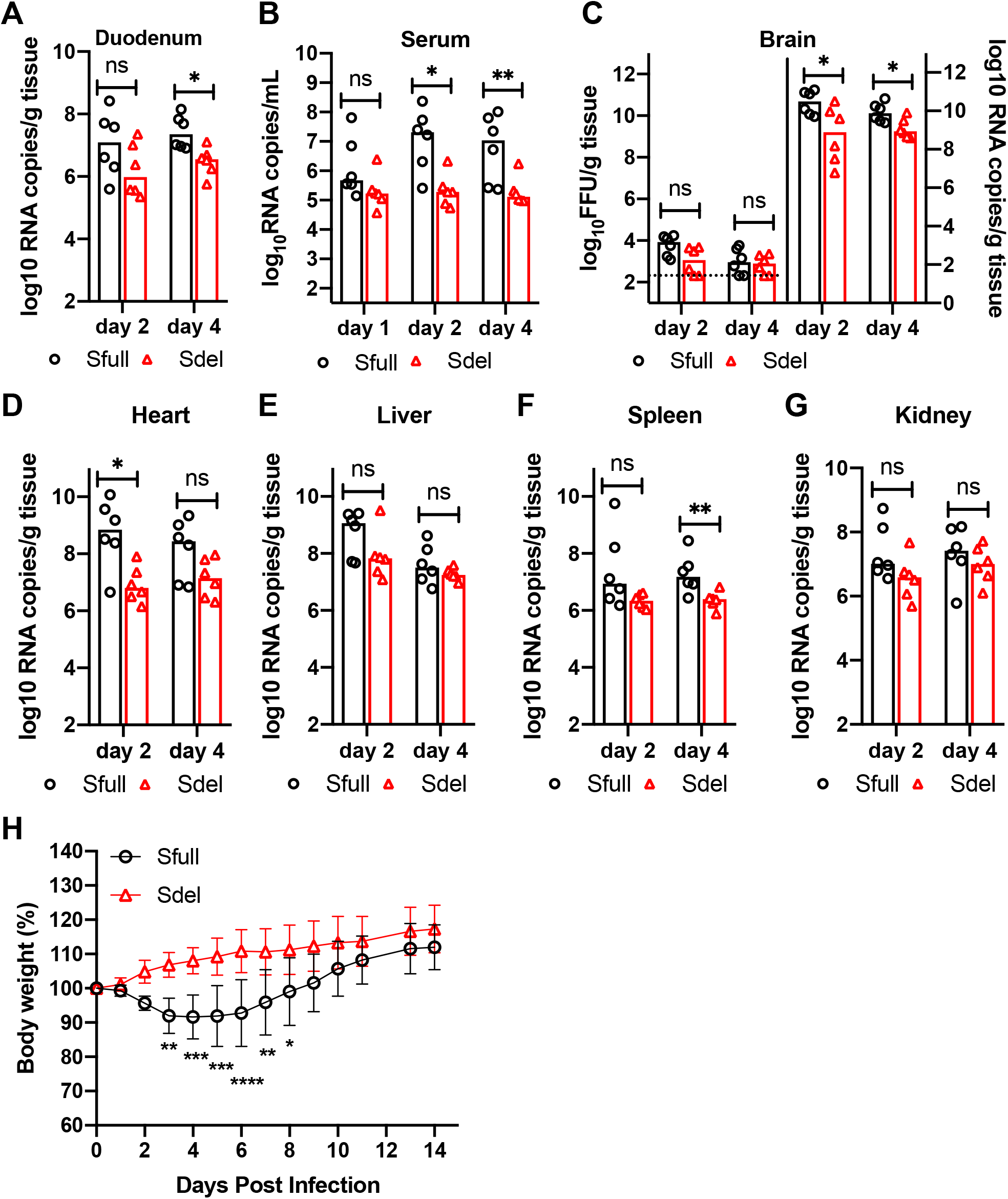
Viral load in different tissues, body weight change and lung histology. **A-G.** 6-8 week-old hamsters were infected intranasally and serum (day 1, 2, 4) and tissues from intestine, brain, heart, liver, spleen, and kidney (day 2, 4) were harvested (n=6 per day). Viral RNAs were extracted for RT-qPCR analysis. The viral load in the brain was also titrated by focus-forming assay. The dashed lines represent the limit of detection by focus-forming assay. Median viral titers: two-tailed Mann–Whitney test *P < 0.05; **P < 0.01; ns, not significant. **H.** Body weight change of hamsters inoculated intranasally with Sfull or Sdel virus. 6-8 week-old hamsters (n=6) were infected with Sfull or Sdel virus and the body weight was measured daily until day 14. Mean body weight: two-way ANOVA with Sidak’s test *P < 0.05; **P < 0.01; ***, P < 0.001; ****P < 0.0001.

## SUPPLEMENTAL TABLE LEGENDS

**Supplementary Table 1. List of genes and scores after MaGeck analysis (see Excel file).** Data was obtained by deep-sequencing of sgRNAs from uninfected or survived cells.

**Supplementary Table 2. sgRNA sequences of genes selected for validation and other editing experiments (see Excel file).**

## Notes

### Competing Interest Statement

The authors have declared no competing interest.

